# Water as a thermal contrast agent for artificial-intelligence-enhanced *in vivo* mid-infrared thermography

**DOI:** 10.64898/2026.07.03.736311

**Authors:** Sixin Xu, Yuanhua Liu, Danyang Xu, Zideng Dai, Wentao Ye, Xingzu Zhan, Feifei Wang

**Affiliations:** Department of Electrical and Computer Engineering, The University of Hong Kong, Hong Kong SAR, China; Materials Innovation Institute for Life Sciences and Energy (MILES), The University of Hong Kong-SIRI, Shenzhen, China; School of Biomedical Engineering, The University of Hong Kong, Hong Kong SAR, China

## Abstract

*In vivo* infrared thermography is limited by the inherently poor spatial resolution at long wavelengths, low contrast, and the lack of biocompatible contrast agents. Here, we present 3–5 μm mid-wave infrared (MWIR) thermography enhanced by an artificial intelligence (AI) network and cold phosphate-buffered saline (PBS) as a thermal contrast agent for noninvasive *in vivo* imaging with high contrast and resolution. MWIR imaging enabled high thermal sensitivity with microscale spatial resolution, strong relative thermal contrast, and facilitated visualization of the subcutaneous vasculature in the human arm, hand, ankle, the femoral artery and vein in rats, and the femoral vessels in mice, with image contrast further enhanced by AI networks. In a 4T1 tumor-bearing mouse model, AI-enhanced MWIR resolved early-stage tumors of ∼2.3 mm and metastases as small as ∼1.7 mm. Using cold PBS as a MWIR thermal contrast agent, we achieved precise tumor boundary visualization and real-time imaging-guided tumor resection. AI-enhanced MWIR offers a promising solution for early diagnosis and improved surgical precision.

## Introduction

Optical infrared imaging in the near infrared (NIR, 0.7–3 μm),^1–7^ mid-wave infrared (MWIR, 3–8 μm),^8–11^ and long-wave infrared (LWIR, 8–14 μm) ^12–14^ spectral windows serves distinct roles in the noncontact, nonionizing assessment of tumors and other diseases. For example, fluorescence imaging in the near-infrared II (NIR-II, 1–3 μm) window enables deep penetration into biological tissues, facilitating the high-contrast, high-resolution imaging of tumors,^15–18^ lymph nodes,^19–21^ immune cells^22–24^ and vasculature^25–27^ in the tumor microenvironment using exogenous probes^28,29^. In particular, imaging at wavelengths greater than 1500 nm in the NIR-II window provides the clearest fluorescence images with minimal tissue autofluorescence,^15,30–33^ but it often utilizes quantum dots or nanoparticles containing toxic elements such as Pb, Cd, and As, raising concerns about biocompatibility and metabolism^34^.

In contrast, infrared thermography derives signals from body thermal radiation, with emissive power following Stefan–Boltzmann’s law (*E* = *εσT*^4^, where *ε* is the emissivity of the emitting surface at a specific wavelength and absolute temperature *T*, and *σ* is the Stefan Boltzmann constant), and has been widely explored in medical applications^35^. Although human tissues exhibit emissivity values of 0.85-0.98 for wavelengths in the range of 2–14 μm,^35^ a very narrow wavelength band (8–12 μm) within the LWIR is generally used in medical thermography, for example, to monitor trauma,^36,37^ infection,^38,39^ inflammation,^40,41^ malignancy-induced temperature increases,^42^ and ischemia-induced temperature decreases^43,44^. Despite these applications, the structural resolving power and sensitivity of LWIR imaging are constrained by weak temperature gradients^45^, spatially variable tissue emissivity^46,47^, ambient thermal noise^48^, and limited spatial resolution.

MWIR thermography emerges as a promising alternative to LWIR imaging, offering theoretically superior spatial resolution due to its shorter wavelengths and deeper tissue penetration attributable to its lower water and tissue absorbance (Fig. 1a-d). Traditionally, it has been believed that water’s absorption in the MWIR window limits its utility for *in vivo* imaging. Fourier transform infrared spectroscopy (FTIR) analysis of deionized water (Fig. 1a) showed strong absorption peaks at 3.0 μm (O-H stretching) and 4.7 μm (a combination of bending and libration modes), between which there is a local minimum in absorption at 3.9 μm. The 2.2-μm spectral band is believed to be the final NIR-II sub-region for one-photon excitation imaging^19^. Although water absorption at 3.9 μm is 1.4 times higher than at 2.2 μm, the attenuation length at 3.9 μm is similar to that at 2.2 μm due to suppressed tissue scattering at longer wavelengths^15^. Tissue absorption in mouse skin, subcutaneous fat, and muscle in the MWIR and LWIR windows is still dominated by water (Fig. 1b-d), while other biomolecules, such as lipids and proteins, contribute only weak, narrow absorption bands. These spectra show an absorption minimum in the 3.9–5.5 μm MWIR window, which is lower than that in the LWIR window. Therefore, MWIR still has potential for *in vivo* imaging in regions with absorption minima.

**Figure 1.**
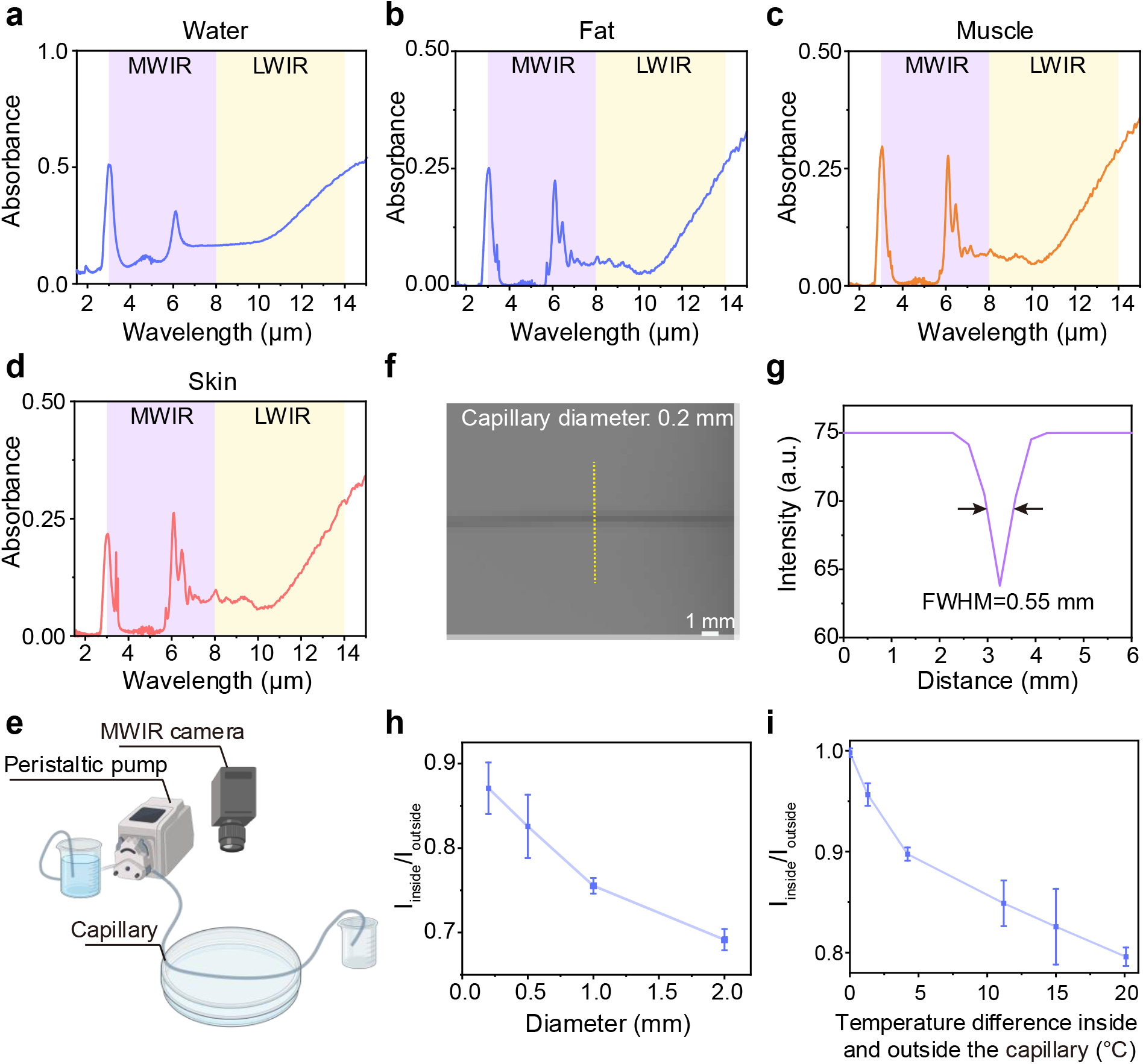
MWIR thermal imaging of water in a capillary tube. Absorbance of (**a**) deionized water, (**b**) fat, (**c**) muscle and (**d**) skin in the MWIR and LWIR windows. (**e**) Schematic diagram of MWIR imaging of a capillary tube placed in a water container. (**f**) MWIR imaging was performed on a capillary tube (inner diameter: 0.2 mm) connected to a peristaltic pump that circulated deionized water at 25 ℃. The tube was immersed in deionized water at 40 ℃, with its top surface level with the water surface in the container. (**g**) Intensity profiles along the dotted lines in **f**. (**h**) Intensity ratios (*I*_inside_/*I*_outside_) between the inside and outside of capillaries with different inner diameters (0.2 mm, 0.5 mm, 1 mm, and 2 mm). The water temperature inside and outside the capillary tube were 25 ℃ and 40 ℃, respectively. (**i**) Intensity ratios for different temperature differences between the inside and outside of the capillary tube. The inner diameter of the capillary tube was 0.5 mm.

Although the absolute thermal signal is stronger in the LWIR range than in the MWIR range, when considering the relative thermal contrast derived from Planck’s law at body temperatures, MWIR imaging can yield higher image contrast than LWIR, as approximately given by:

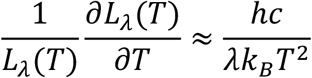

where *L*_*λ*_(*T*) is radiance, *T* is absolute temperature, *λ* is wavelength, *ℎ* is Planck’s constant, *c* is speed of light, *k*_*B*_ is Boltzmann constant.

MWIR thermography has shown potential in noninvasive surface tumor assessment, and its higher resolution than LWIR aids in delineating boundaries associated with malignancy. For example, it can distinguish between benign and malignant tumors based on their behavior during dynamic cooling and temperature recovery^49^, and detect melanoma metastases larger than 15 mm in diameter with 95% sensitivity and 100% specificity^50^. However, identifying small tumors using MWIR is challenging because thermal signals from the body and ambient noise reduce imaging contrast.

Here, we present artificial neural network-enhanced MWIR thermography for visualization of vasculature in humans, rats, and mice, as well as early-stage tumor lesions or small metastases *in vivo*, with high contrast and resolution. It enabled longitudinally imaging of the process of tumor growth and proliferation, and monitoring the relative temperature of the tumor compared to surrounding healthy tissues. Cold phosphate-buffered saline (PBS) has been explored as an MWIR thermal contrast agent for tumor labeling, enabling precise identification of tumor boundaries and MWIR imaging-guided tumor resection.

## Results

### MWIR thermal imaging of water in a capillary tube

To evaluate the feasibility of using water as a thermal contrast agent, we performed 3–5 µm MWIR imaging of capillaries that were filled with deionized water circulated by a peristaltic pump at an average flow velocity of 10 cm/s to maintain the temperature as constant as possible and immersed in a temperature-controlled water bath (Fig. 1e). The temperatures of the circulating water in the capillary and the water in the water bath were 25 ℃ and 40 ℃, respectively, creating a temperature difference of 15 ℃. When a capillary with an inner diameter of 0.2 mm was used, we observed thermal contrast between the water inside and outside the capillary in the MWIR image (Fig. 1f), with a full width at half maximum (FWHM) of 0.55 mm (Fig. 1g). As the inner diameter of the capillary increased from 0.2 mm to 2 mm, the image contrast increased, and the signal ratio between the inside and outside of the tube decreased from 0.87 to 0.69 (a lower ratio indicates higher contrast; Fig. 1h; Supplementary Fig. 1). Furthermore, at a fixed tube diameter of 0.5 mm, increasing the temperature difference between the inside and outside of the tube from 0 ℃ to 20 ℃ enhanced the imaging contrast and decreased the inside-to-outside intensity ratio from 0.998 to 0.795 (Fig. 1i, Supplementary Fig. 2), confirming that the temperature difference influences the MWIR imaging contrast. These results demonstrate the high sensitivity of MWIR imaging in detecting micrometer-scale temperature differences and validate the use of water as a thermal contrast agent in MWIR thermography.

### Enhanced vascular visualization in MWIR thermography via a deep learning network

Non-invasive imaging of microvascular structure is critical for vascular disease diagnosis and minimally invasive surgical guidance. We evaluated the vascular visualization capabilities of MWIR thermography in humans, rats and mice, and applied CycleGAN,^51^ a deep learning framework for image-to-image translation that does not require paired training data, to enhance MWIR image resolution and contrast (Methods).

We first imaged the human arm, hand and ankle in the MWIR (3–5 µm) and LWIR (8–14 µm) windows (Fig. 2). To avoid potential hardware bias among different LWIR cameras, we performed additional side-by-side comparisons using three LWIR cameras (Luster, Hikvision, Hti; Supplementary Table 1) under identical experimental conditions. The LWIR thermography showed the temperature distribution in the human arm, hand and ankle with low contrast and without clear structural detail (Fig. 2c,g,k; Supplementary Fig. 3). In contrast, MWIR thermography revealed the subcutaneous vascular network with high clarity (Fig. 2a,e,i). This contrast arises from the higher temperature of blood compared with the surrounding tissues. The FWHMs of the blood vessels in the human arm, hand, and ankle in the original MWIR images were smaller than those in the LWIR images recorded by the Luster Cobra8000 (0.68 cm vs. 1.06 cm, 0.66 cm vs. 1.06 cm, 0.65 cm vs. 1.48 cm; Fig. 2d,h,l). Meanwhile, the signal-to-background ratios (SBRs) of the vessels in the MWIR images of the arm, hand, and ankle were higher than those in the LWIR images: 6.5 vs. 6.1, 7.7 vs. 4.8, and 9.7 vs. 8.3, respectively. After neural-network processing of the MWIR images, we achieved high SBRs of 30.3, 34.6 and 40.2 for the vessels in the human arm, hand and ankle, respectively, which were 4.7, 4.5 and 4.1 times higher than those of the raw MWIR images, significantly improving image contrast and clarity.

**Figure 2.**
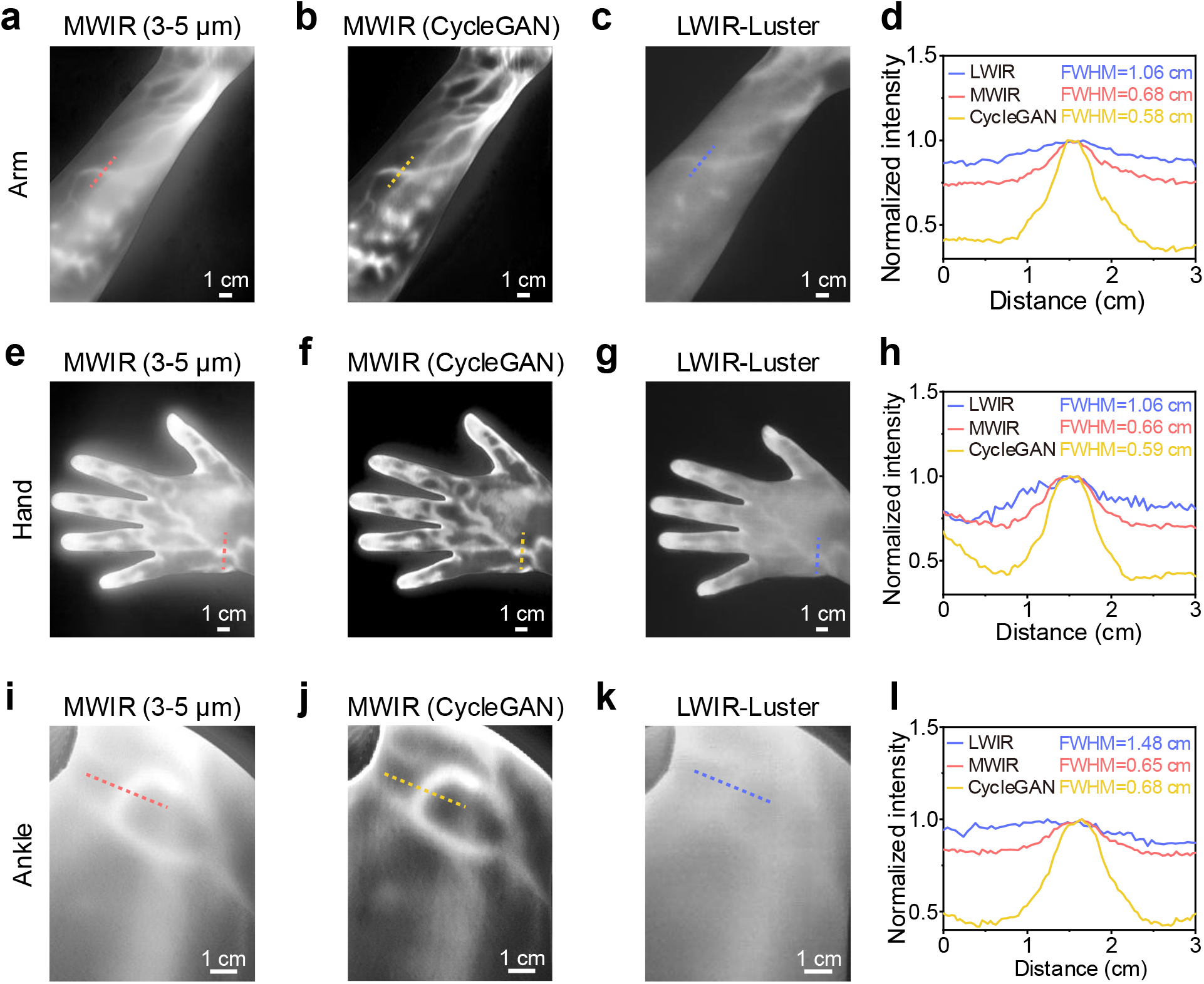
Human blood vessel imaging in the MWIR and LWIR windows. (**a**) MWIR image, (**b**) CycleGAN-enhanced MWIR image, and (**c**) LWIR image of a human arm. (**d**) Normalized intensity profiles along the dotted lines in **a, b** and **c**. (**e**) MWIR image, (**f**) CycleGAN-enhanced MWIR image, and (**g**) LWIR image of a human hand. (**h**) Normalized intensity profiles along the dotted lines in **e, f** and **g**. (**i**) MWIR image, (**j**) CycleGAN-enhanced MWIR image, and (**k**) LWIR image of a human ankle. (**l**) Normalized intensity profiles along the dotted lines in **i, j** and **k**.

We then performed MWIR thermography in rats and observed the femoral artery and vein in the hindlimb (Fig. 3a-e, Supplementary Fig. 4a), which are difficult to identify clearly in LWIR images recorded by the Hikvision and Hti cameras (Supplementary Fig. 4d,e and 5a). Although the artery and vein can be resolved in the LWIR images recorded by the Luster Cobra8000, the MWIR image shows higher contrast between artery and vein than the LWIR image (1.13 vs. 1.05; Supplementary Fig. 4a,c). The FWHM of the femoral artery was ∼1.32 mm in the MWIR images and increased to 2.47 mm in the LWIR images recorded by the Luster Cobra8000 (Supplementary Fig. 4f). The SBR of artery increased from 1.15 in the raw MWIR image to 1.54 after CycleGAN processing (Supplementary Fig. 4a,b). In the magnified MWIR image of another rat hindlimb, the FWHM of the femoral artery was ∼1.83 mm and decreased to 1.49 mm after CycleGAN processing, while the SBR increased from initial 1.06 to 1.26 (Fig. 3c,d), confirming improved resolution and contrast. In addition to enhanced contrast and resolution in blood vessel imaging, the abdominal imaging contrast was also improved (Fig. 3a,b), revealing the temperature distribution at higher resolution, which was not achievable with LWIR thermography (Supplementary Fig. 5a).

**Figure 3.**
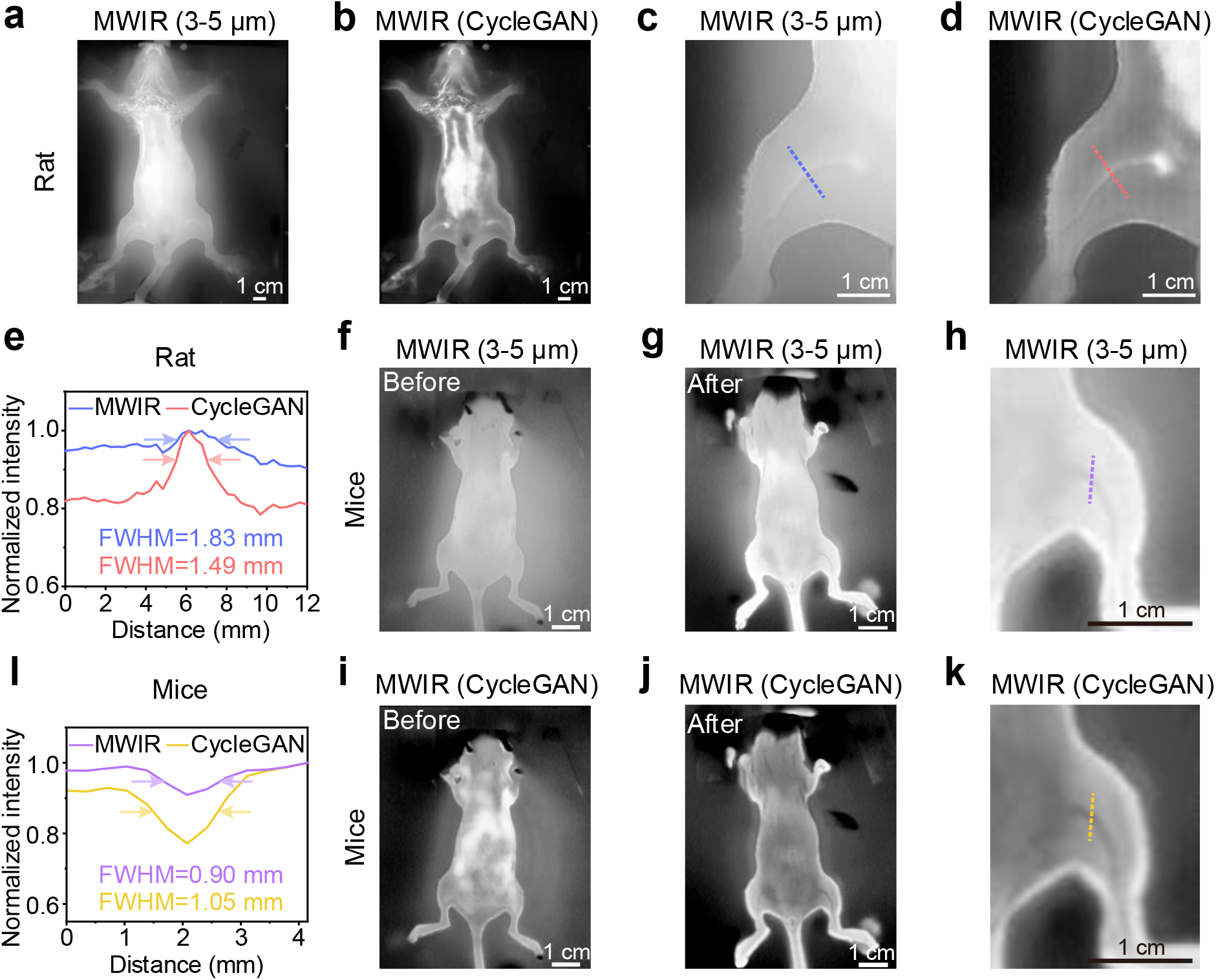
MWIR imaging of blood vessels in rats and mice. (**a**) MWIR image and (**b**) CycleGAN-enhanced MWIR image of an entire rat. (**c**) MWIR image and (**d**) CycleGAN-enhanced MWIR image of a rat hindlimb. (**e**) Normalized intensity profiles along the dotted lines in **c** and **d**, shown in blue and red, respectively. (**f**,**g**,**h**) MWIR images and (**i**,**j**,**k**) CycleGAN-enhanced MWIR images of a mouse (**f**,**i**) before, and (**g**,**h**,**j**,**k**) after being immersed in 39 ℃ water for 5 seconds. (**l**) Normalized intensity profiles along the dotted lines in **h** and **k**, shown in purple and yellow, respectively.

Although MWIR thermography enabled visualization of human blood vessels and the rat femoral artery and vein, we did not resolve blood vessels in mice in either the raw or CycleGAN-processed images (Fig. 3f,i). Consistent with the findings in rats, the mouse femoral blood vessel was also not identifiable in visible and LWIR images (Supplementary Fig. 5b). However, after immersing the mouse’s hindlimb in 39 ℃ warm water for 5 seconds, the femoral blood vessel became visible in the MWIR window (Fig. 3g,h). Although the FWHM of the femoral blood vessel increased from 0.90 mm to 1.05 mm (Fig. 3j), the SBR decreased from 0.92 to 0.75 after neural network processing, facilitating better contrast (Fig. 3h,k,l). We note that the SBR is less than 1 because the temperature of the blood vessel is lower than that of the surrounding tissues. Under this condition, the smaller the SBR value, the higher the contrast. We further imaged the dynamically thermo-regulated mice using one MWIR camera and three LWIR cameras (Luster, Hikvision, Hti; Supplementary Fig. 6). For LWIR imaging, the mouse femoral veins could be resolved with the Luster Cobra8000 camera (Supplementary Fig. 6c) but remained invisible with the Hti and Hikvision cameras (Supplementary Fig. 6d,e). However, the MWIR provides better contrast (0.77 vs. 0.78) and higher resolution (0.67 mm vs. 1.04 mm) than LWIR imaging with the Luster Cobra8000 camera (Supplementary Fig. 6a,c,f). The SBR of femoral blood vessels in the MWIR window was reduced from 0.77 to 0.45 using CycleGAN, enabling better visualization.

### Longitudinal monitoring of tumor growth via artificial intelligence (AI)-enhanced MWIR thermography

Noninvasive, high-contrast, real-time visualization of tumors *in vivo* is critical for early diagnosis and surgical precision^52–55^. Assessing tumor temperature is a promising approach, as tumor temperatures can be either lower or higher than those of surrounding normal tissues, depending on factors such as metabolic activity, vascularization, and progression stage^56^. For example, hypothermic tumors frequently exhibit temperatures below body temperature due to reduced metabolic heat generation in hypoxic regions and impaired blood perfusion, which limits heat exchange^57^. In contrast, hyperthermic tumors, such as those in pancreatic ductal adenocarcinoma (PDAC), show intrinsic temperature increases driven by heightened lipid metabolism^58^. Both hypothermia and hyperthermia have the potential to serve as early biomarkers of tumor progression, enabling non-invasive tracking via MWIR imaging.

We established a 4T1 tumor-bearing mouse model to explore the feasibility of longitudinal monitoring of early tumor growth using CycleGAN-enhanced MWIR thermography and to compare MWIR with LWIR imaging acquired using three commercial LWIR cameras (Fig. 4a). After 4T1 cell inoculation, tumor-bearing mice were monitored longitudinally for five days in the visible (Fig. 4b), LWIR (Hikvision, Hti, and Luster Cobra8000 cameras; Fig. 4c and Supplementary Fig. 7-9) and MWIR (Fig. 4d) windows. On day 1, MWIR imaging resolved the injection needle puncture site on the mouse abdomen (FWHM = 0.69 mm) (Fig. 4d,f, Day 1), whereas none of the three LWIR cameras detected any corresponding structure (Fig. 4c and Supplementary Fig. 7-9). MWIR thermography enabled identification of tumor on day 2 and longitudinal monitoring of its growth over subsequent days (Fig. 4d,f). In contrast, LWIR images recorded by the Luster Cobra8000 camera showed no detectable tumor on days 2-3 and only blurry, low-contrast outlines on days 4-5 (Fig. 4c; Supplementary Fig. 7-9). The Hikvision LWIR camera resolved the tumor on day 5 with very low contrast, while the Hti LWIR camera did not detect it on any of the imaging days (Supplementary Fig. 7-9). All these results demonstrate that MWIR provides better imaging resolution and sensitivity than LWIR.

**Figure 4.**
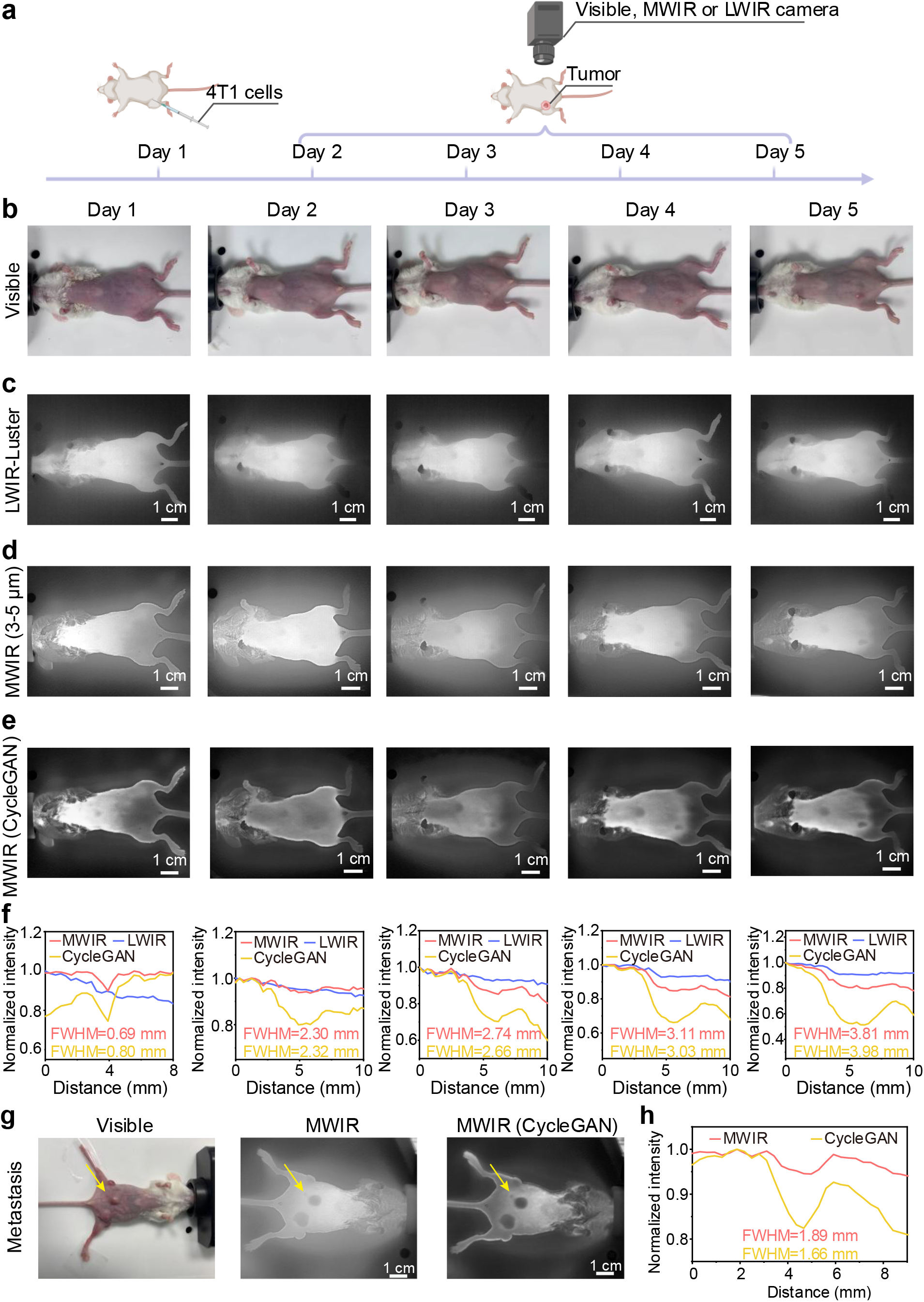
Longitudinal MWIR imaging of 4T1 tumor-bearing mice. (**a**) Tumor inoculation and imaging schedule. 4T1 cells were inoculated on day 1. Imaging in the visible, MWIR, and LWIR windows was performed on days 1-5. (**b**) Visible images, (**c**) LWIR images recorded by the Luster Cobra8000 camera, (**d**) MWIR images and (**e**) CycleGAN-enhanced MWIR images of a 4T1 tumor-bearing mouse on days 1-5. (**f**) Normalized intensity profiles along the dotted lines crossing the tumor, which were marked in Supplementary Fig. 7c, d, e, are shown in blue, red and yellow, respectively. (**g**) Visible, MWIR, and CycleGAN-enhanced MWIR imaging of a mouse bearing three major 4T1 tumors and a small metastasis. The arrows indicate the small metastasis in the MWIR and CycleGAN-enhanced MWIR images. (**h**) Normalized intensity profiles along the dotted lines crossing the metastasis, which were marked in Supplementary Fig. 10a,b, are shown in red and yellow, respectively.

Tumor-to-normal tissue (T/NT) ratio can be further enhanced via CycleGAN processing, facilitating the recognition of tumor boundaries (Fig. 4e). On days 2-5, the tumor FWHM showed no obvious change in the processed MWIR images compared with the raw images (2.30 mm vs. 2.32 mm, 2.74 mm vs. 2.66 mm, 3.11 mm vs. 3.03 mm, 3.81 mm vs. 3.98 mm), but the T/NT ratio decreased to 0.86-, 0.86-, 0.83- and 0.75-fold, respectively (Fig. 4d-f). The decrease in the T/NT ratio enabled better contrast because the tumor temperature was lower than that of surrounding tissue (Fig. 4d,e).

In a mouse bearing three major 4T1 tumors, we observed small metastases with the FWHM of 1.89 mm in the MWIR image (Fig. 4g,h; Supplementary Fig. 10). The FWHM of metastases decreased to 1.66 mm, and the contrast of metastases can be further enhanced by 1.13-fold using the CycleGAN model, facilitating early tumor diagnosis and treatment evaluation. The temperature of the 4T1 tumor was lower than that of the surrounding tissues, potentially due to the decreased tumor cell metabolic activity in the hypoxic core region^56^. AI-enhanced MWIR thermography provides a high-resolution, high-sensitivity method for tumor temperature mapping, allowing for exploration of the relationship between tumor progression and temperature changes.

### Water as a MWIR thermal contrast agent for precise tumor resection

We then explored the use of cold PBS as a low-temperature thermal contrast agent in MWIR imaging to enhance tumor visibility. PBS-enhanced MWIR is achieved with MWIR-guided injection, preserves the MWIR-defined tumor outline and transiently increases T/NT contrast for surgical guidance. Specifically, 50 µL of 4 ℃ phosphate-buffered saline (PBS) solution was injected intratumorally into a 4T1 tumor-bearing mouse as a low-temperature MWIR thermal contrast agent for tumor labeling (Fig. 5a). MWIR thermography was performed in the 3–5 µm window during intratumoral PBS injection, revealing a rapid decrease in the tumor signal. The SBR decreased from 0.92 to 0.13 after PBS injection (with SBR < 1 because the tumor temperature was lower than that of the surrounding normal tissues), resulting in a sevenfold increase in imaging contrast (Fig. 5b). The tumor signal gradually returned to its pre-injection level within 200 seconds post-injection (Fig. 5c). To prolong the observation window, the injection can be repeated, or the boundary can be marked before the thermal contrast dissipates for real-time guidance. Processing the images with the CycleGAN neural network (Fig. 5d) further improved tumor contrast in images acquired before and after PBS injection, demonstrating the feasibility of using water as a thermal contrast agent for MWIR thermography to selectively enhance tumor contrast via temperature regulation.

**Figure 5.**
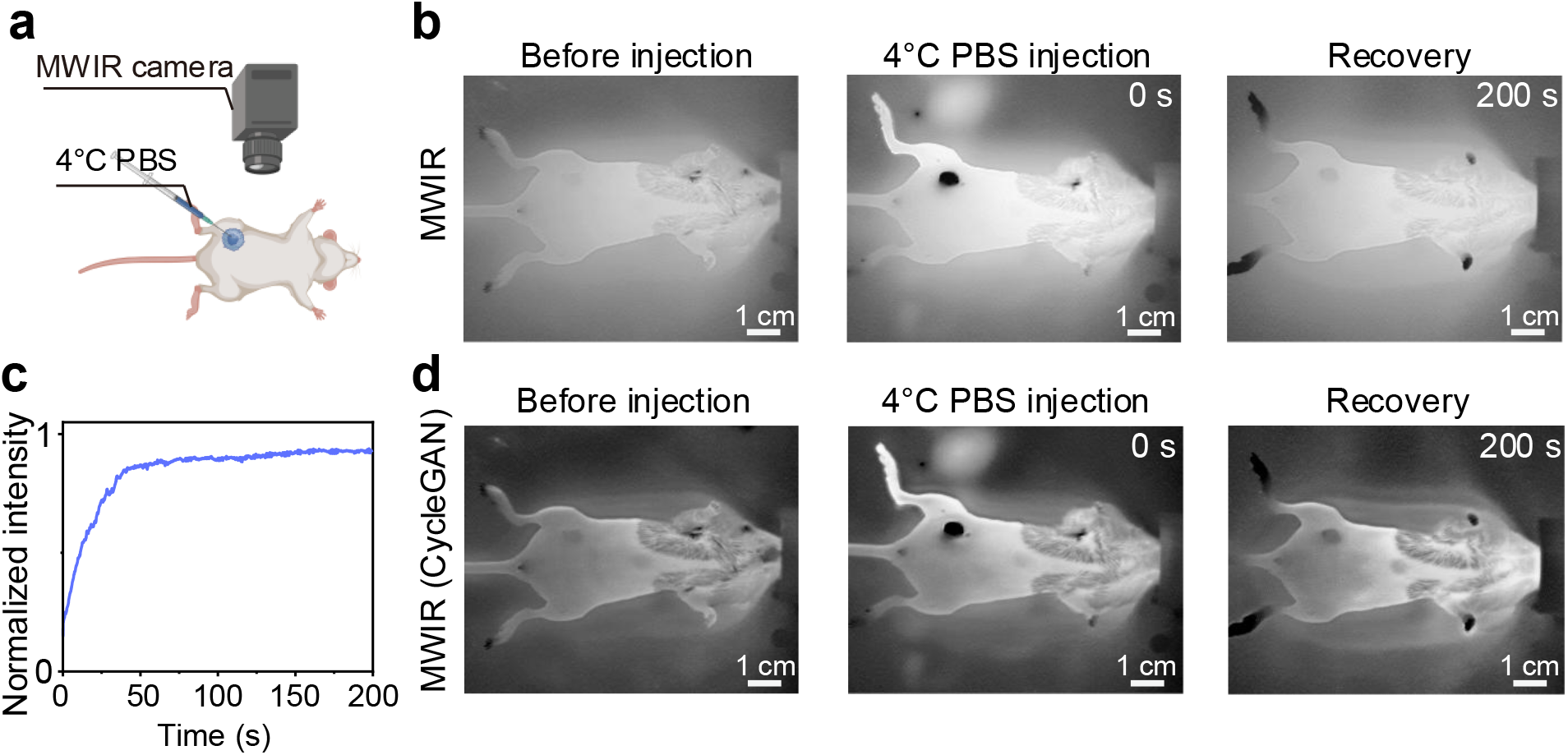
Water as a thermal contrast agent for MWIR imaging. (**a**) Schematic diagram of MWIR imaging of a 4T1 tumor-bearing mouse using water as a thermal contrast agent injected intratumorally. (**b**) MWIR imaging of a 4T1 tumor-bearing mouse before and at various time points after intratumoral injection of 4 ℃ PBS. (**c**) Changes in tumor intensity in a 4T1 tumor-bearing mouse after intratumoral injection of 4 ℃ PBS. (**d**) CycleGAN-processed MWIR images correspond to those shown in **b**.

Next, we performed MWIR thermography-guided 4T1 tumor resection surgery in the 3–5 µm spectral window using cold water as the thermal contrast agent (Fig. 6a). The frame rate of MWIR imaging was 50 frames per second (fps). After intratumoral injection of 4 ℃ PBS as a low-temperature thermal contrast agent, MWIR imaging clearly revealed the tumor boundary with high contrast and precision (Step 2, Fig. 6b), which could be further enhanced by CycleGAN processing (Fig. 6c). Cross-sectional line profiles across the tumor showed a significant decrease in tumor signal intensity after PBS injection (Fig. 6d), providing much stronger contrast for surgical navigation than in label-free MWIR thermography-guided surgery (Supplementary Fig. 11). Using low-temperature water as a thermal contrast agent for MWIR imaging guidance, complete tumor resection was successfully achieved (Step 3, Fig. 6b,c), providing higher contrast and resolution than LWIR (Fig. 6e) and visible imaging (Fig. 6f). To further validate that the tumor region identified by MWIR imaging corresponded to actual tumor tissue rather than image-processing artifacts, we performed H&E staining of the tumor after MWIR-guided resection. Histological examination confirmed the presence of tumor tissue in the resected lesion, and the tumor boundary was consistent with the region identified by MWIR imaging before surgery (Supplementary Fig. 12).

**Figure 6.**
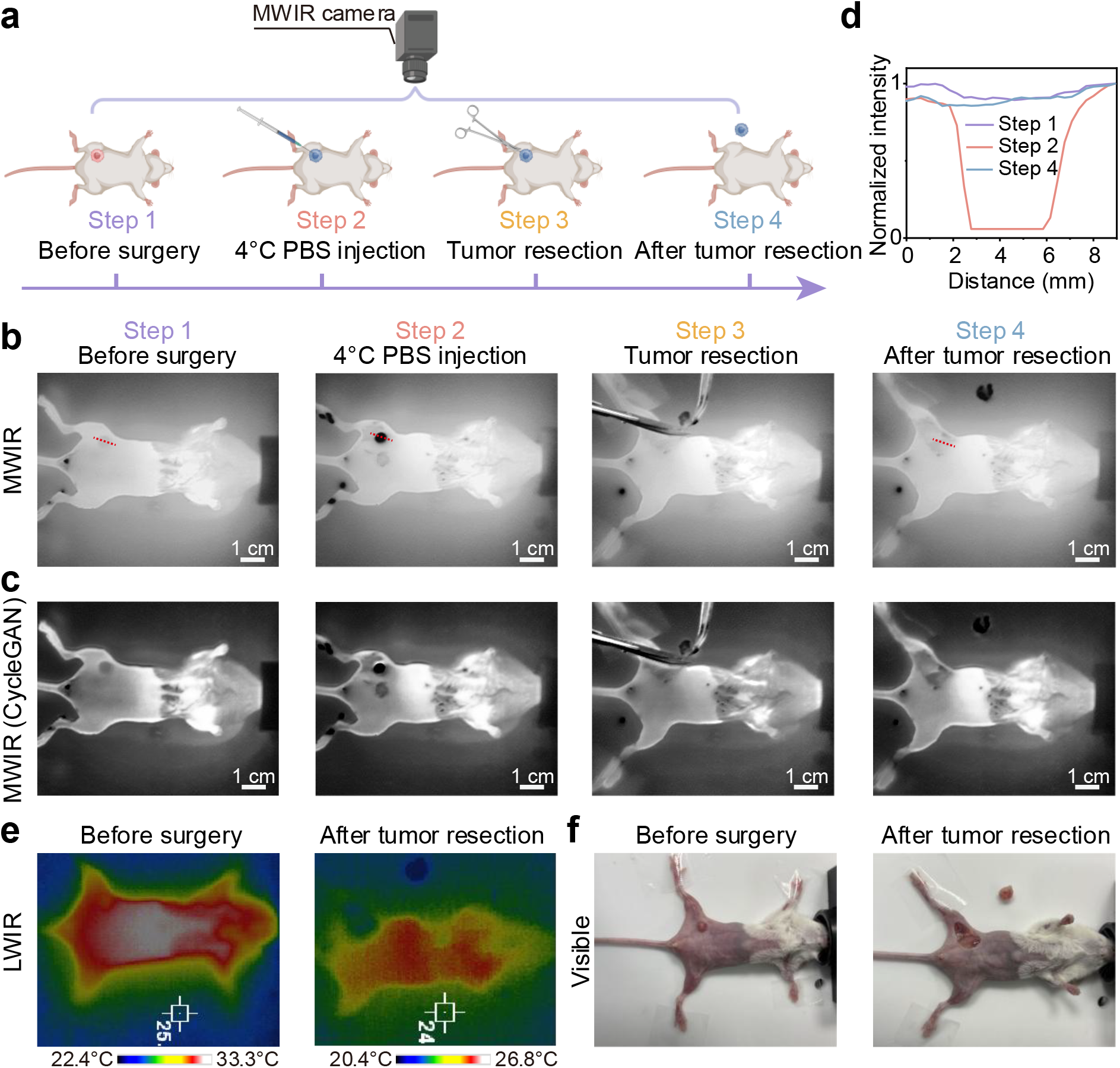
Water as a MWIR thermal contrast agent for precise tumor resection. (**a**) Experimental schedule for MWIR-guided surgical removal of a 4T1 tumor from a mouse. Tumor resection guided by (**b**) MWIR and (**c**) CycleGAN-enhanced MWIR imaging, using 4 ℃ PBS as a thermal contrast agent. (**d**) Normalized intensity profiles along the dotted lines in the Step 1 (before injection of 4 ℃ PBS), Step 2 (after 4 ℃ PBS injection), and Step 4 (after tumor resection) images in **b**. (**e**) LWIR and (**f**) visible imaging of a 4T1 tumor-bearing mouse before injection of 4 ℃ PBS and after tumor removal.

## Discussion

Accurate identification of tumor margins with high image clarity and contrast is a key requirement for precise image-guided tumor resection that minimizes damage to healthy tissue. This work demonstrated the feasibility of MWIR thermography for *in vivo* imaging and using water as a MWIR natural thermal contrast agent for precise tumor resection. High-sensitivity MWIR imaging with ∼0.55 mm resolution was achieved by imaging capillaries with micrometer-scale temperature gradients. With the enhancement of a deep-learning-based method (CycleGAN network), *in vivo* label-free MWIR thermography of the human subcutaneous vasculature network and the rat femoral artery and vein was achieved with high contrast and spatial resolution, overcoming the technical limitations of traditional LWIR imaging. By leveraging the natural temperature gradients arising from tumor metabolic activity^57^, we observed early-stage microtumors (diameter: ∼2.3 mm) and metastases (diameter: ∼1.7 mm) with high sensitivity. Based on the approximate expression for relative thermal contrast derived from Planck’s law at body temperatures (*T* = 310 K), the relative thermal contrast is estimated to be 3.74%/K at 4 μm in the MWIR window and 1.36%/K at 11 μm in the LWIR region, indicating an ∼ 2.75-fold improvement. Further direct comparisons with MWIR and LWIR cameras at similar levels confirm that the observed advantages (e.g., higher resolution and contrast) are not merely due to hardware differences but arise from the fundamental wavelength-dependent benefit of the MWIR window. Another advantage of MWIR over LWIR is its lower absorption by water and tissue, facilitating deeper tissue penetration.

The use of low-temperature PBS as a safe, low-cost, exogenous MWIR thermal contrast agent significantly enhanced tumor contrast relative to surrounding healthy tissues, enabling precise visualization of tumor boundaries and real-time MWIR thermography-guided surgery. However, the contrast induced by cold water gradually diminished within 200 seconds, which may raise concerns for long-duration surgery. To address this issue, artificial learning-based automatic segmentation can be performed on tumor lesions during the initial high-contrast phase, leveraging early-frame MWIR images to accurately identify tumor boundaries. Segmented tumor areas can then be highlighted with pseudo-color overlays to facilitate intraoperative visualization^65^.

Deep learning methods provided an effective means of improving the imaging performance of MWIR thermography, yielding higher contrast and resolution. Expanding and diversifying the training dataset can further mitigate overfitting and improve generalizability across diverse tissue types. A real-time neural-network implementation will enable intraoperative image enhancement and is critical for tracking rapid physiological dynamics. AI-enhanced MWIR thermography offers a promising approach to disease diagnosis and precision surgery.

In clinical practice, LWIR thermography has been utilized to detect temperature differences as potential biomarkers in skin and breast cancer screening^66^ and applied to assess vascular diseases^67^ and inflammation monitoring^68^. To explore the applicability and potential of MWIR in humans, we imaged vascular structures in both the MWIR and LWIR windows, including subcutaneous veins in the forearm (depth, ∼1-2 mm)^69^, dorsal hand veins (depth, ∼0.5-2 mm)^70^, and ankle veins (depth, ∼3-5 mm)^71^ (Fig. 2). These vessels were more clearly visualized in the MWIR images than in the LWIR images, with sharper boundaries and higher relative thermal contrast arising from blood temperature differences. These results highlight the potential of MWIR for non-invasive, high-resolution thermal imaging in humans, even without contrast agents. Although deep-tissue penetration remains limited, this does not preclude its application in fields such as dermatology, screening for peripheral vascular diseases, and assessment of tumor margins in superficial layers or in exposed tumors during surgery.

## Methods

### Materials

PBS buffer, RPMI 1640 medium and fetal bovine serum (FBS) were purchased from Gibco (USA). The peristaltic pump was purchased from Leirong Co., Ltd (China).

### Characterization

Nicolet iS50 Fourier transform infrared spectrometer (ThermoFisher, USA) was applied to collect Fourier transform infrared spectra.

### MWIR and LWIR imaging system

MWIR thermography was performed using a Cobra5000 camera (Luster, China) and a 3 μm long-pass filter was used. LWIR images were recorded by multiple infrared thermal cameras (Cobra8000, Luster, China; HM-TPK20, Hikvision, China; HT-A2, Hti-Xintai Instrument, China). Supplementary Table 1 provides detailed specifications for each camera. The MWIR Luster Cobra5000 camera and the LWIR Luster Cobra8000 camera were fitted with fixed-focal-length 25 mm lenses.

### Establishment of 4T1 tumor-bearing mice

All animal experiments were approved by the Materials Innovation Institute for Life Sciences and Energy (MILES), The University of Hong Kong-SIRI. Four-week-old SD male rats and six-week-old BALB/c female mice were purchased from Charles River. The 4T1 cells were cultured in RPMI 1640 medium supplemented with 10 % (v/v) FBS at 37 ℃ in a 5 % (v/v) CO_2_ humidified atmosphere. 4T1 cells (1×10^7^ cells) suspended in 100 μL PBS were subcutaneously injected into each mouse to establish a 4T1 tumor-bearing mouse model. Mice were selected randomly from cages for experiments.

### MWIR thermography and image-guided tumor removal *in vivo*

MWIR thermography of rats and mice as well as image-guided tumor removal surgery in 4T1 tumor-bearing mice, were performed under a rodent anesthesia machine using 2 L/min O_2_ gas mixed with 3 % isoflurane.

### Deep learning algorithm training

The training code for CycleGAN was adapted from the original framework^72^. Specifically, the U-Net architecture was used as the generator to transfer low-quality images to high-quality images, while PatchGAN was the discriminator architecture for evaluating the performance of the generator. The network structure and training hyperparameters were kept consistent with those used in our previous work ^51^. We trained the model for a total of 300 epochs, with an initial learning rate of 0.0002 for the first 100 epochs, followed by a linear decay to zero over the subsequent 200 epochs. Model performance was evaluated every 50 epochs. The entire deep learning algorithm was implemented using PyTorch 2.7.1 with Python 3.12.2. All training and inference steps were performed on an NVIDIA RTX 4090 GPU with 24 GB of memory.

### Statistical and data analysis

All analyses were conducted using OriginPro (2021). All images were analyzed using ImageJ (Version 1.53c) for intensity extraction and quantitative analysis.

## Data availability

All data that support the findings of this study are available in the main text or the supplementary materials.

## Acknowledgments

This study was supported by National Natural Science Foundation of China (Project No. T2522030), Early Career Scheme (RGC No. 27204623) and General Research Fund (RGC No. 17212424) from the Research Grants Council of Hong Kong SAR, the JC STEM Lab of Nanoscience and Nanomedicine funded by The Hong Kong Jockey Club Charities Trust, NSFC/RGC Collaborative Research Scheme (CRS_HKU703/25) and start-up funding from Materials Innovation Institute for Life Sciences and Energy (MILES), HKU-SIRI in Shenzhen.

## Author contributions

F.W. and S.X. conceived and designed the experiments. S.X. performed MWIR thermography. Y.L. and X.Z. processed MWIR images using a CycleGAN model. S.X. and F.W. wrote the manuscript. All authors contributed to the general discussion and revision of the manuscript.

## Materials & Correspondence

Correspondence and requests for materials should be addressed to F.W.

