## Supplementary Materials for "Water as a thermal contrast agent for artificial-intelligence-enhanced *in vivo* mid-infrared thermography"

### SUPPLEMENTARY FIGURES

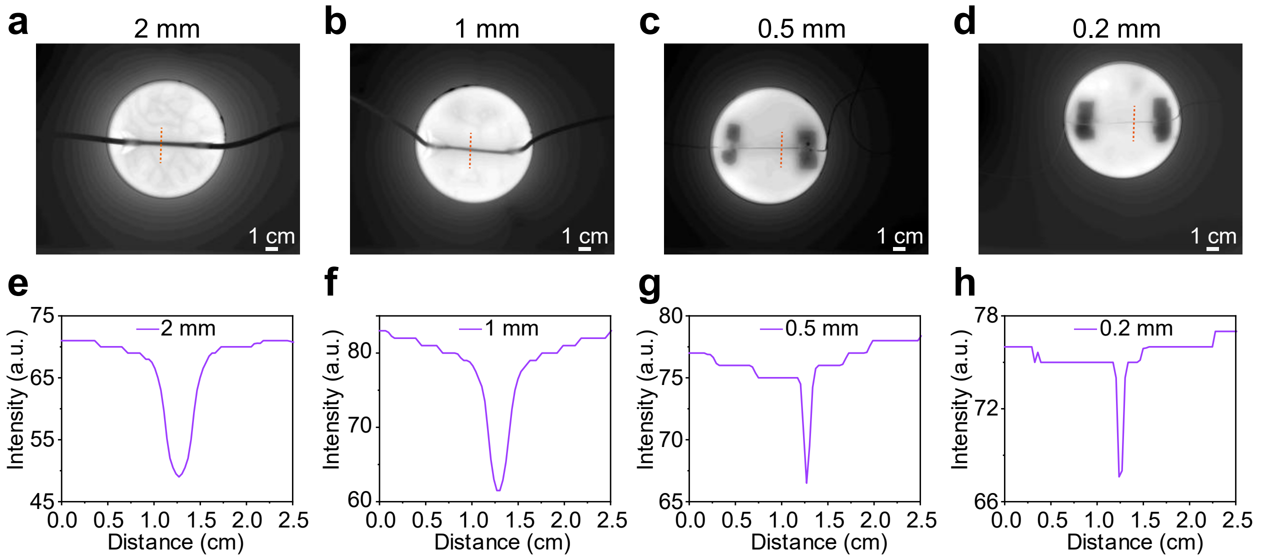


**Supplementary Figure 1. MWIR imaging and intensity profiles of capillary tubes with different inner diameters immersed in a water.** (**a-d**) MWIR imaging of capillary tubes with different inner diameters (2 mm, 1 mm, 0.5 mm, and 0.2 mm) immersed in a water container. The water temperature inside the tubes and outside the tubes was 25 °C and 40 °C, respectively. (**e-h**) Intensity profiles along the dotted lines in **a-d**.

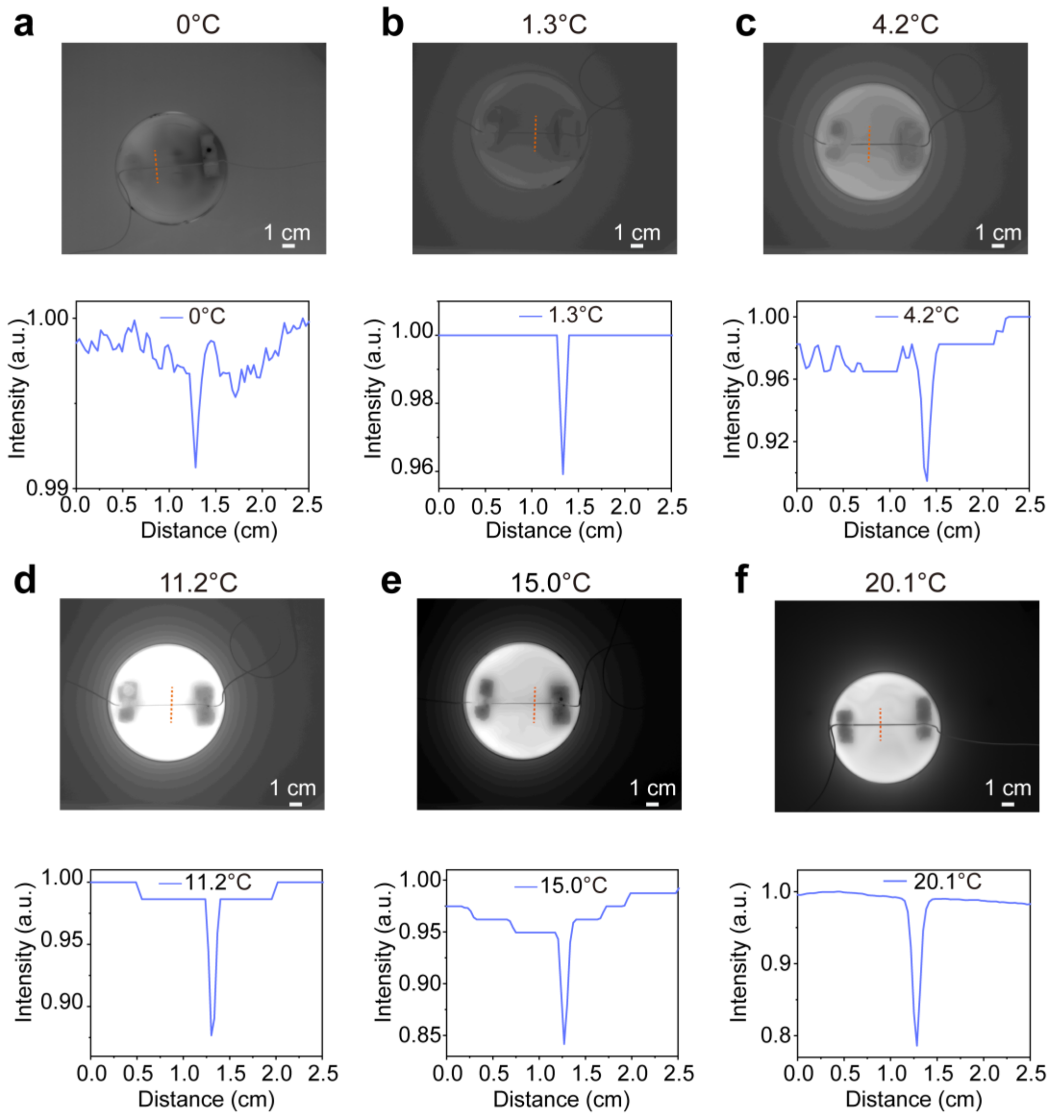


**Supplementary Figure 2. MWIR imaging and intensity profiles of the capillary tube under different temperature differences between the inside and outside water.** **(a-f)** MWIR imaging and intensity profiles along the dotted lines in MWIR images with different temperature differences between the inside and outside the capillary (0 ℃, 1.3 ℃, 4.2 ℃, 11.2 ℃, 15.0 ℃, 20.1 ℃). The inner diameter of the capillary tube is 0.5 mm.

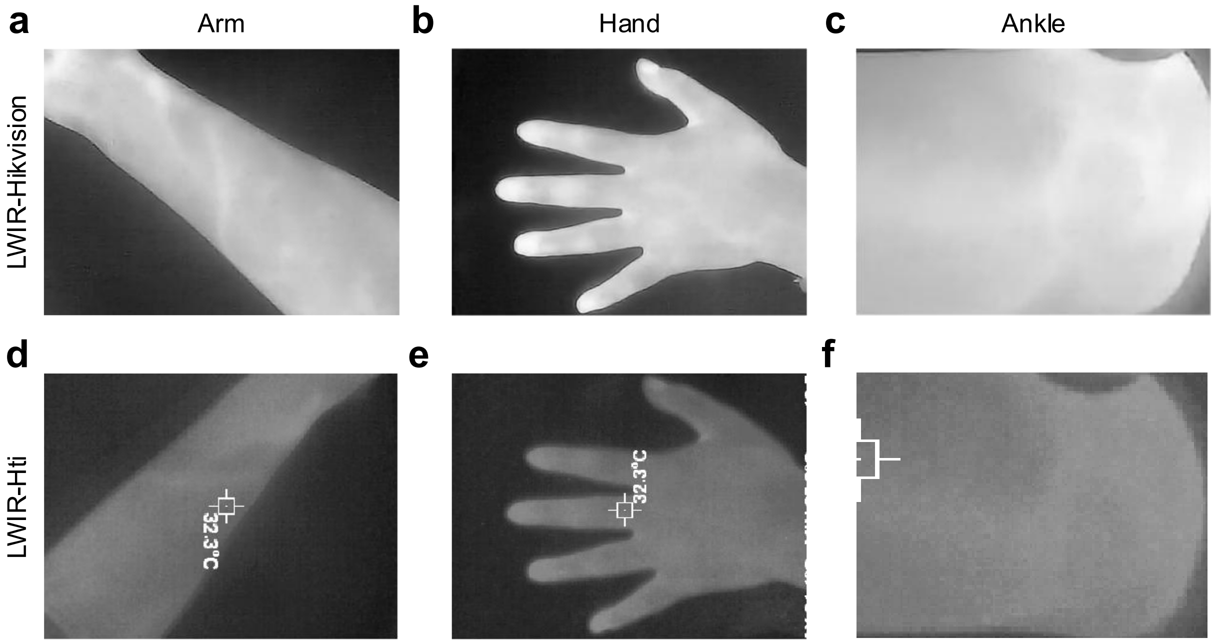


**Supplementary Figure 3. LWIR imaging of blood vessels in the human arm, hand and ankle.** The LWIR images of the (**a**) human arm, (**b**) hand and (**c**) ankle were recorded by a Hikvision camera. The LWIR images of a (**d**) human arm, (**e**) hand and (**f**) ankle were recorded by an Hti camera. The corresponding MWIR images are shown in Fig. 2.

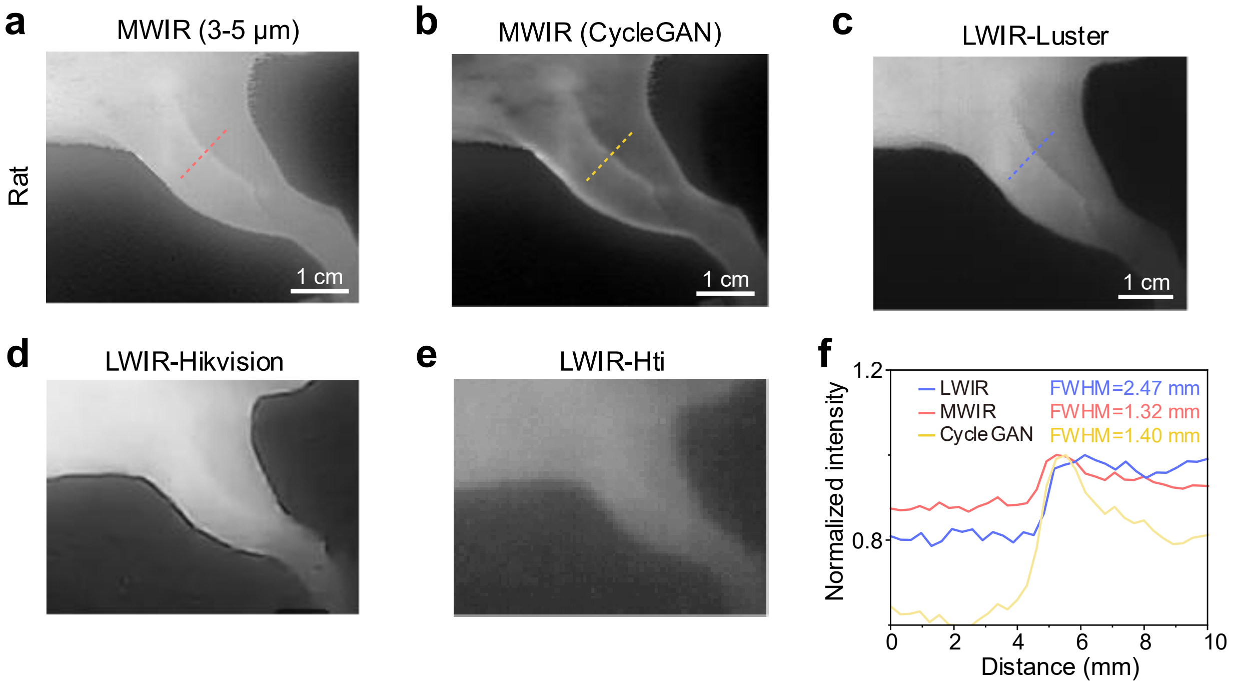


**Supplementary Figure 4. MWIR and LWIR imaging of blood vessels in rats.** (**a**) MWIR image and (**b**) CycleGAN-processed MWIR image of a rat hindlimb. LWIR images of a rat hindlimb acquired with (**c**) the Luster Cobra8000 camera, (**d**) the Hikvision camera, and (**e**) the Hti camera. (**f**) Intensity profile along the dotted lines in **a**, **b** and **c**.

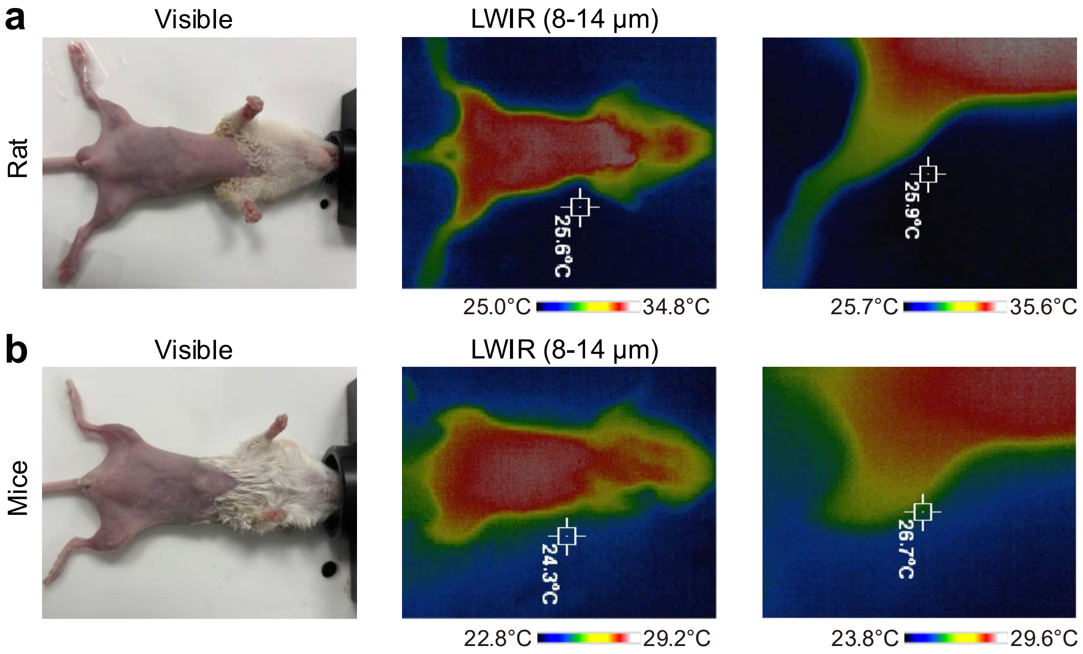


**Supplementary Figure 5. Visible and LWIR imaging of a rat and a mouse.** (**a**) Visible image, LWIR image, and local zoomed LWIR image of a rat. (**b**) Visible image, LWIR image, and local zoomed LWIR image of a mouse.

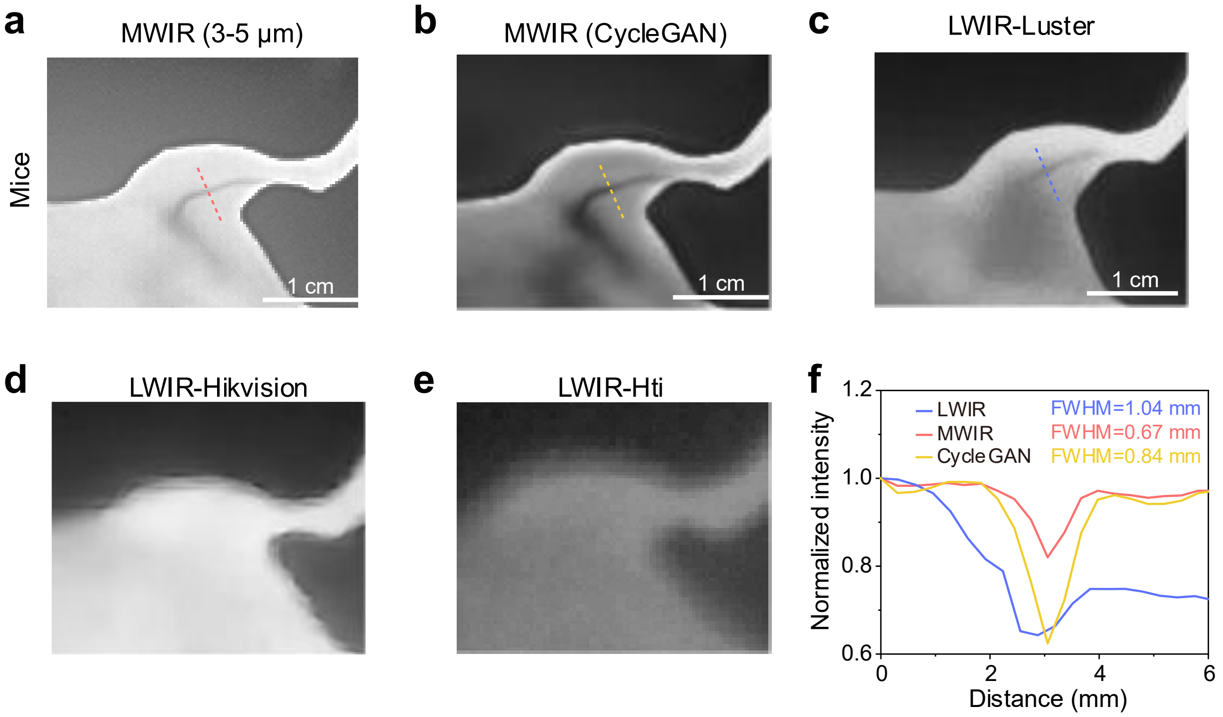


**Supplementary Figure 6. MWIR and LWIR imaging of blood vessels in mice.** (**a**) MWIR image and (**b**) CycleGAN-processed MWIR image of a mouse hindlimb. LWIR images of a mouse hindlimb acquired with (**c**) the Luster Cobra8000 camera, (**d**) the Hikvision camera, and (**e**) the Hti camera. (**f**) Intensity profile along the dotted lines in **a**, **b** and **c**. The mouse was immersed in 39 °C water for 5 seconds before MWIR and LWIR imaging.

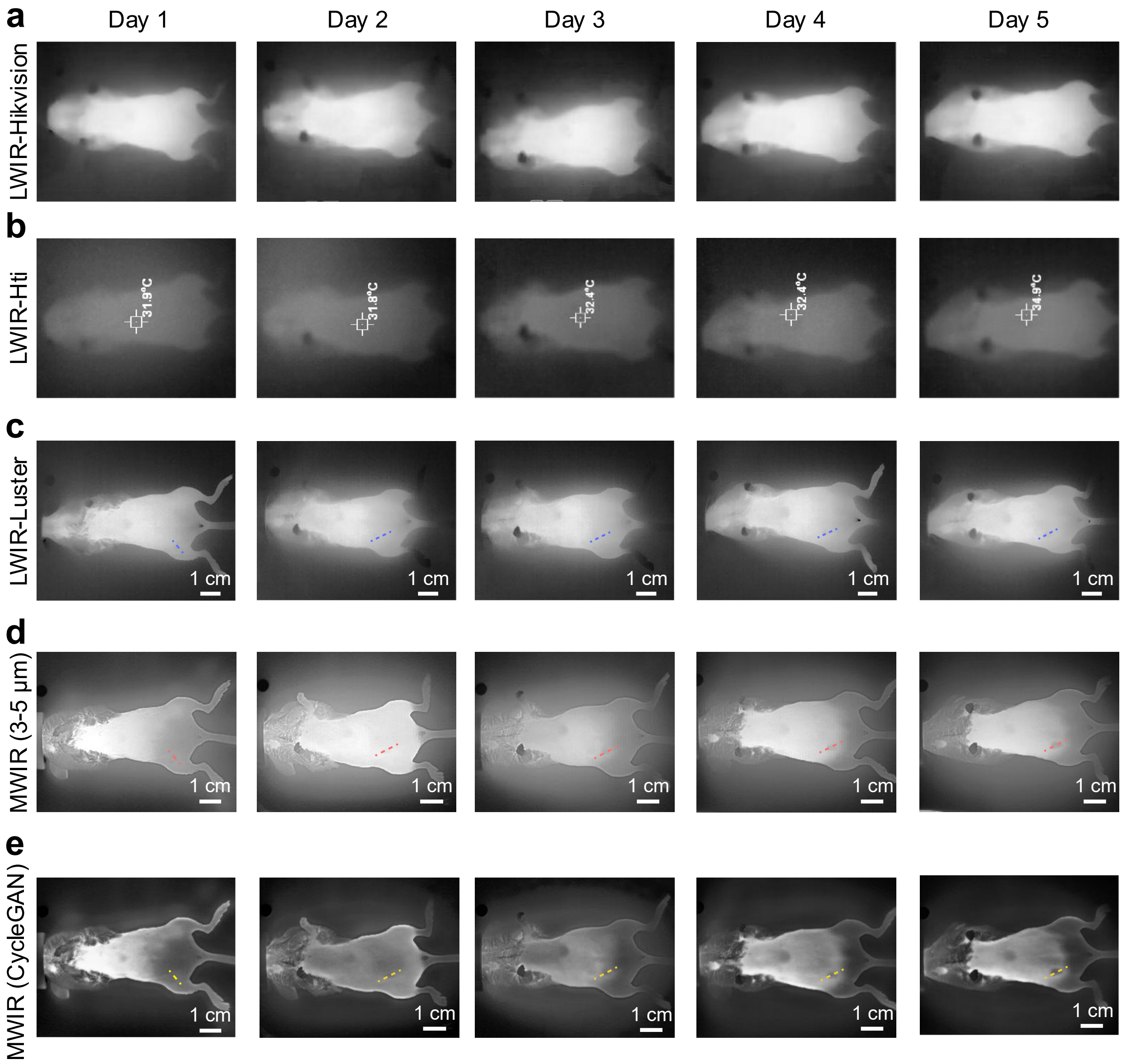


**Supplementary Figure 7. Longitudinal LWIR imaging of the first 4T1 tumor-bearing mouse.** LWIR images of a 4T1 tumor-bearing mouse over days 1-5 were recorded using (**a**) the Hikvision camera, (**b**) the Hti camera, and (**c**) the Luster Cobra8000 camera. The corresponding (**d**) MWIR and (**e**) CycleGAN-enhanced MWIR images of the same 4T1 tumor-bearing mouse over days 1-5. 4T1 cells were inoculated on day 1. The images shown in **c**-**e** are the same as those in Figure 4c-e.

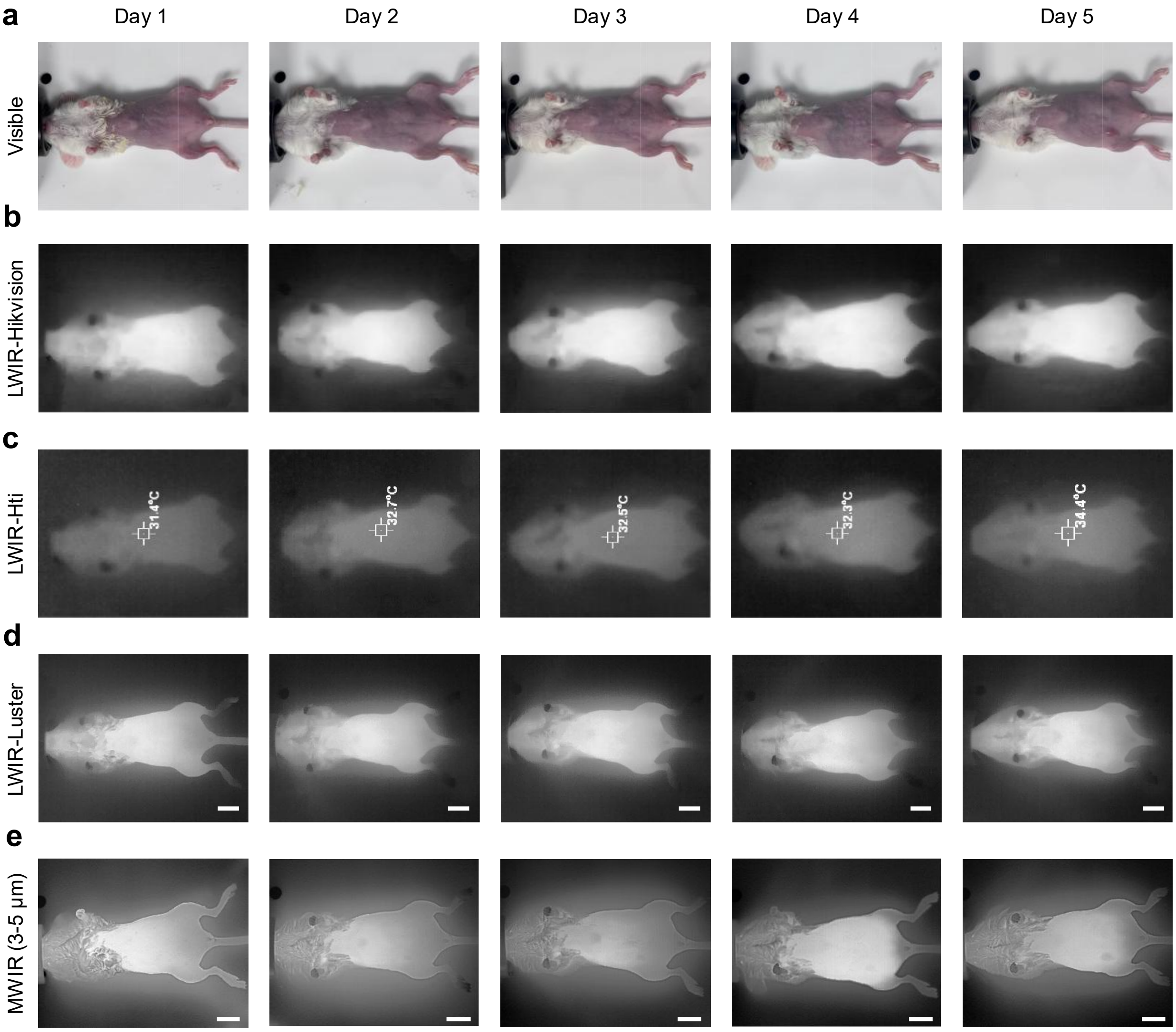


**Supplementary Figure 8. Longitudinal MWIR imaging of the second 4T1 tumor-bearing mouse.** (**a**) Visible image, (**b**) LWIR image (Hikvision), (**c**) LWIR image (Hti), (**d**) LWIR image (Luster Cobra8000), and (**e**) MWIR image of a 4T1 tumor-bearing mouse on days 1-5. Scale bar = 1 cm.

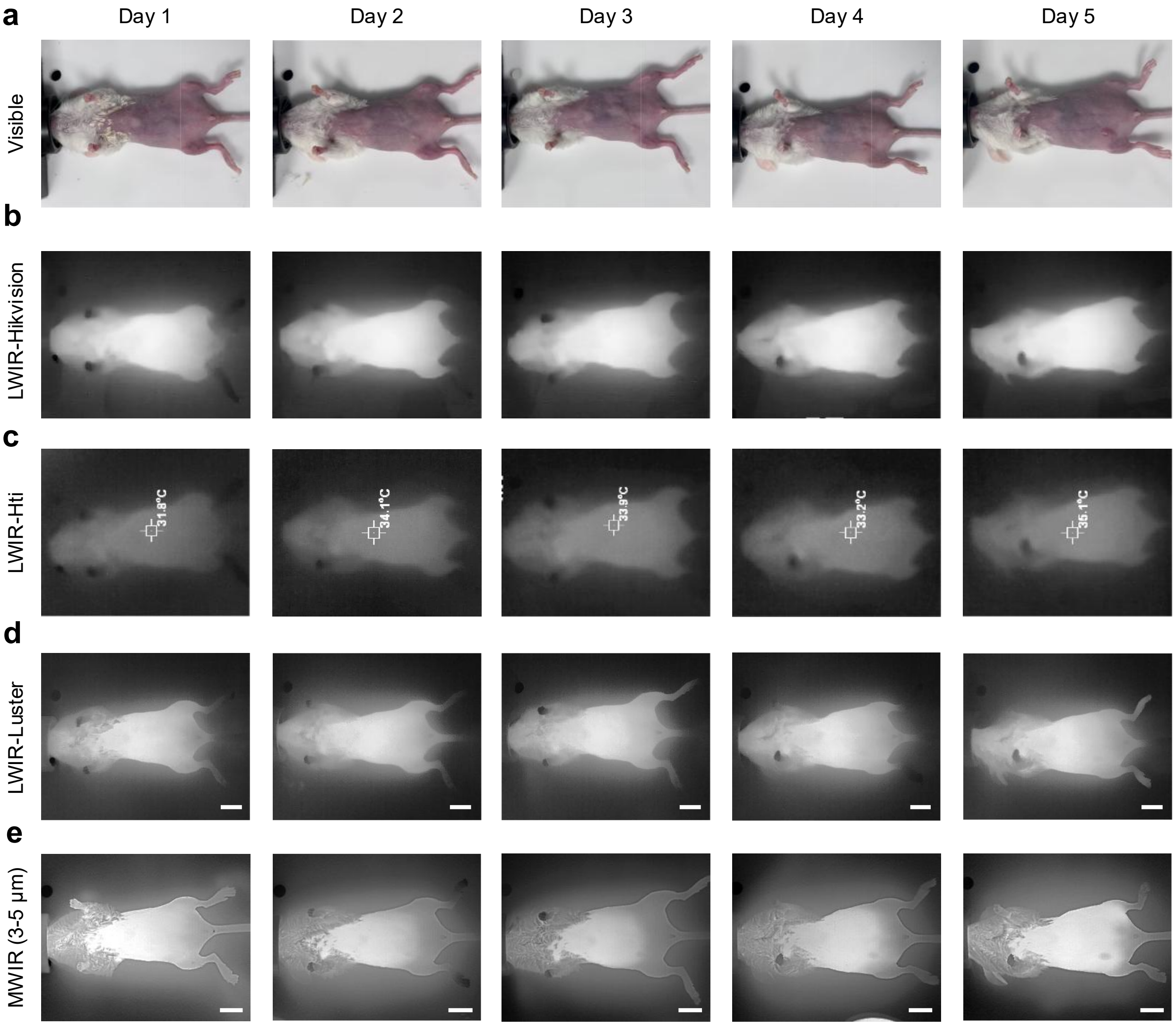


**Supplementary Figure 9. Longitudinal MWIR imaging of the third 4T1 tumor-bearing mouse.** (**a**) Visible image, (**b**) LWIR image (Hikvision), (**c**) LWIR image (Hti), (**d**) LWIR image (Luster Cobra8000), and (**e**) MWIR image of a 4T1 tumor-bearing mouse on days 1-5. Scale bar = 1 cm.

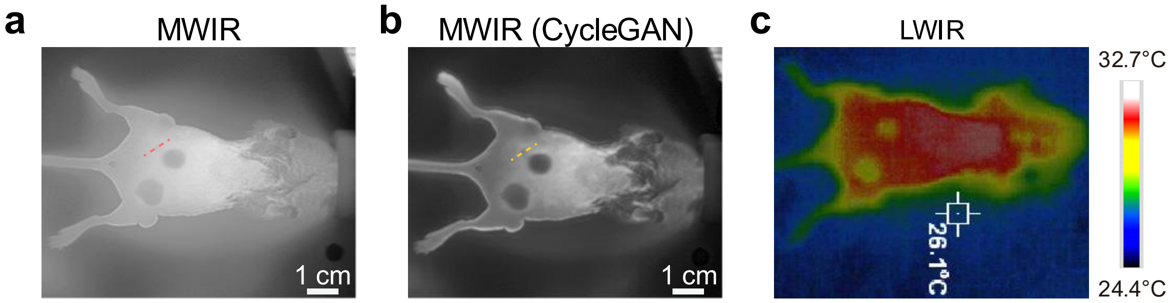


**Supplementary Figure 10. MWIR imaging of metastasis in 4T1 tumor-bearing mice.** (**a**) MWIR, (**b**) CycleGAN-enhanced MWIR and (**c**) LWIR imaging of a mouse bearing three major 4T1 tumors and a small metastasis. Dotted lines cross the tumor in image a,b. Images of a and b are the same as those in Figure 4g. To avoid obscuring the tumors in Figure 4g, dotted lines were only added to these images.

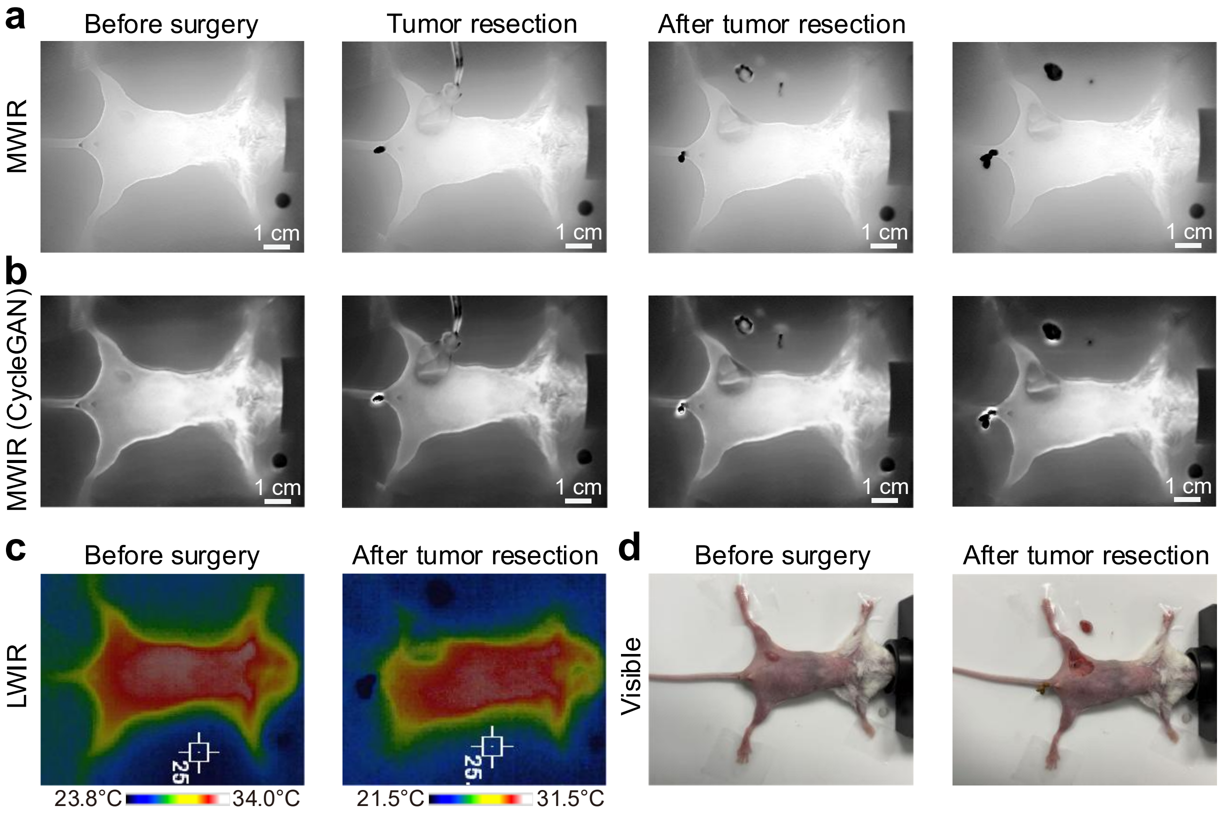


**Supplementary Figure 11. MWIR thermography-guided tumor resection.** Tumor resection guided by (**a**) MWIR and (**b**) CycleGAN-enhanced MWIR imaging. (**c**) LWIR and (**d**) visible imaging of a 4T1 tumor-bearing mouse before and after tumor resection. 4T1 cells were inoculated 4 days before imaging.

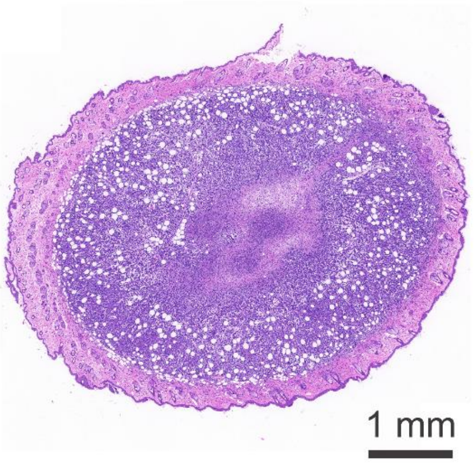


**Supplementary Figure 12. H&E staining and histological examination of the tumor tissue resected under MWIR imaging guidance.**

### Supplementary Table 1. Parameters of MWIR and LWIR cameras

| | MWIR | LWIR | | |
|---|---|---|---|---|
| Brand | Luster | Luster | Hikvision | Hti-Xintai |
| Camera Model | Cobra5000 | Cobra8000 | HM-TPK20 | HT-A2 |
| Corresponding band | 1.5 -5 μm | 8 -14 μm | 8 -14 μm | 8-14 μm |
| Resolution | 640×512 | 640×512 | 256×192 | 320×240 |
| Pixel size | 15 μm | 12 μm | 12 μm | |
| NETD | < 20 mK | < 50 mK | < 50 mK | < 70 mK |

### Supplementary Table 2. The FWHMs of the blood vessels in humans, rats, and mice imaged in MWIR and LWIR windows.

| | | MWIR (Luster, Cobra5000) | LWIR (Luster, Cobra8000) |
|---|---|---|---|
| Human | Arm | 0.68 cm | 1.06 cm |
| | Hand | 0.66 cm | 1.06 cm |
| | Ankle | 0.65 cm | 1.48 cm |
| Rat | Femoral artery | 1.32 mm | 2.47 mm |
| Mouse | Femoral vein | 0.67 mm | 1.04 mm |

### Supplementary Table 3. The SBRs of the blood vessels in humans, rats, and mice imaged in MWIR and LWIR windows.

| | | MWIR (Luster, Cobra5000) | LWIR (Luster, Cobra8000) |
|---|---|---|---|
| Human | Arm | 6.5 | 6.1 |
| | Hand | 7.7 | 4.8 |
| | Ankle | 9.7 | 8.3 |
| Rat | Femoral artery | 1.15 | 1.14 |
| Mouse | Femoral vein | 0.77 | 0.78 |
